# Marker-based watershed transform method for fully automatic mandibular segmentation from low-dose CBCT images

**DOI:** 10.1101/397166

**Authors:** Yi Fan, Richard Beare, Harold Matthews, Paul Schneider, Nicky Kilpatrick, John Clement, Peter Claes, Anthony Penington, Christopher Adamson

## Introduction

Three-dimensional mandibular models are useful for planning maxillofacial surgery and orthodontic treatment.^1,2^ In studies of growth, mandibular models are important for assessing morphological changes over time.^3,4^ Such models are typically obtained from conventional computed tomography (CT), using high radiation dose to capture fine detail of the bony structure. Cone beam computed tomography (CBCT) shows promise for oral and craniofacial imaging applications due to lower radiation dose, lower cost and shorter acquisition time compared to CT. However, CBCT images have lower contrast and higher levels of noise than conventional CT, making mandible segmentation a challenging task.^5^

Segmenting a 3D mandible is typically done ‘interactively’ in computer software on a case by case basis. Threshold-based algorithms or morphological operations are commonly used first for the separation of bony structures from soft tissues.^6–8^ Then manual work is needed to separate the mandible from the cranial base and the maxilla because the algorithms cannot distinguish between different facial bones with similar intensity values. We refer to this combination of computerized operations and manual editing as ‘interactive’ segmentation. Specific issues in mandible segmentation include intercuspation occlusion and low contrast of condyles relative to surrounding structures. Intercuspation leads to connection between upper and lower teeth while low contrast leads to difficult to define boundaries on the condyles. These require slice-by-slice based manual editing, which is tedious, time-consuming and operator-dependent as it produces slightly different outlines after continuous interventions.^6^

A robust automated mandible segmentation approach is thus desirable. In other applications, simple automated methods based on voxel intensity or edge intensity can be very effective.^9,10^ However, the mandible is not the only bone structure in CBCT images of the head, and the intensity of bone varies considerably. More sophisticated methods which incorporate prior information about the expected shapes and position of objects to be segmented are required. To date only four publications, using statistical shape models,^11,12^ multiatlas label registration^13^ or machine learning,^14^ have been proposed to automate the mandibular segmentation from CBCT images. These approaches are either computationally expensive or require collection of large amounts of manually segmented mandibles as training data, which may be impractical in clinical situations.

The watershed method is a classic, computationally simple technique for object segmentation in images. The original grayscale image can be regarded as a topographic relief, with brightness/intensity corresponding to altitude, and thus identifying watershed lines provides a method for segmenting an image into separate spatial regions.^15,16^ The original grayscale image is transformed into a ‘height map’ which emphasizes discontinuities in image intensity (such as occur at object boundaries) and dampens continuous regions (such as homogeneous intensity region within the tissue). The marker-based watershed transform dilates, or floods, from markers that are provided to the algorithm. The number of markers determines the number of regions that will be created by the watershed transform.^17^ The watershed markers can be manually or automatically set. The marker-based watershed transform has been successfully used to segment breast lesions on ultrasound^18^ and lymphoma in sequential CT images^19^ but never been used to segment mandibles from CBCT images.

In this article, we propose and validate an automatic approach for segmenting mandibles from low-dose CBCT using a marker-based watershed transform. We fully automate the segmentation by automated watershed marker placement using image registration. Segmentation accuracy of the proposed automated method is assessed by comparing outcomes with a well-accepted interactive segmentation method described in the literature.^1,2^

## Methods and materials

### Image data

CBCT images were obtained from 21 adolescent subjects with a mean age of 13.68 years (SD:1.27; 9 males) from an orthodontic clinic, where images had previously been obtained for clinical indications. Images were taken using an i-Cat machine (Imaging Sciences International, PA, USA) with a 16 × 22cm field of view and an isotropic voxel size of 0.5mm. Patients were instructed to bite into maximum intercuspation during scanning. Ethics approval to use the images was obtained from (Hidden Content).

### Overview of the marker-based watershed mandible segmentation

In a CBCT image, the mandible has typically high intensity and therefore is brighter than its surrounding tissue (muscles or air) with a marked drop in intensity at the boundaries. The height map is constructed to enhance these boundaries and suppress homogeneous regions. Here the height map was generated by transforming the CBCT image into gradient image of itself using the Derivative of Gaussian (Full Width at Half Maximum 1mm) kernel, which highlighted boundaries of sharply changing intensity in the original image. Initially, two sets of watershed markers were placed on the gradient image, one set within the mandible and the other set in the rest of the image. The watershed transform floods the gradient image by dilating the markers simultaneously until colliding at watershed lines, estimating the mandible boundary. Illustration of the marker-based watershed transform method was shown in Figure 1. This method was used in both generating template data and automating segmentation of the mandible from novel images.

**Figure 1.**
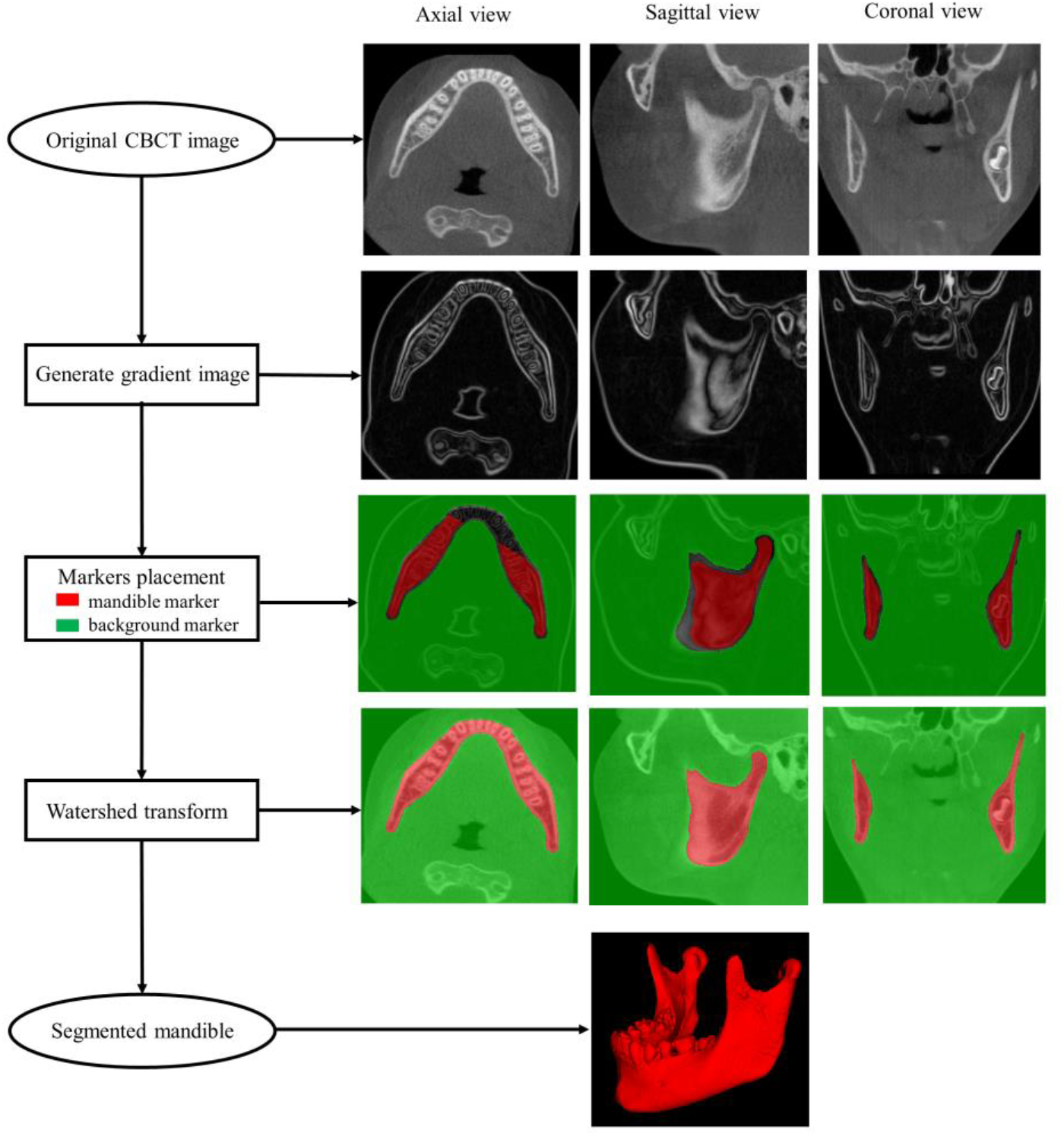
Illustration of the marker-based watershed transform method. The original image is transformed into the gradient image which highlights boundaries of sharply changing intensity in the original image. The mandible marker (in red) and the background marker (in green) are placed within the mandible and at the rest of the structures, separately. The watershed transform floods the gradient image by dilating the markers simultaneously until colliding at watershed lines, estimating the mandible boundary. The segmented mandible is reconstructed in below. The pipeline is demonstrated in axial (column 1) sagittal (column 2) and coronal (column 3) views.

### Template construction

An image of a 12.65 years old male patient was selected to create the template data comprised of a CBCT image and associated mandible and background markers that can later be propagated onto the novel image. The mandible and background markers were defined semi-automatically on the template image by applying the watershed method described above to manually drawn markers (lines or circles that were drawn unambiguously within or without the mandible as shown in Figure 2). The watershed transform segmented the mandible, and the remainder of the image was labelled as background. These segmentations were then eroded by 1 voxel to form the markers that have an unlabeled gap where the expected mandible boundary location resides. The dental crowns were manually removed from the mandible marker because the eruption stages and the number of the teeth could be different between the template image and the test images and the teeth were not of primary interest. The final mandible marker of the template image was displayed in red and with the background marker overlaid in green as shown in the marker placement row in Figure 1.

**Figure 2.**
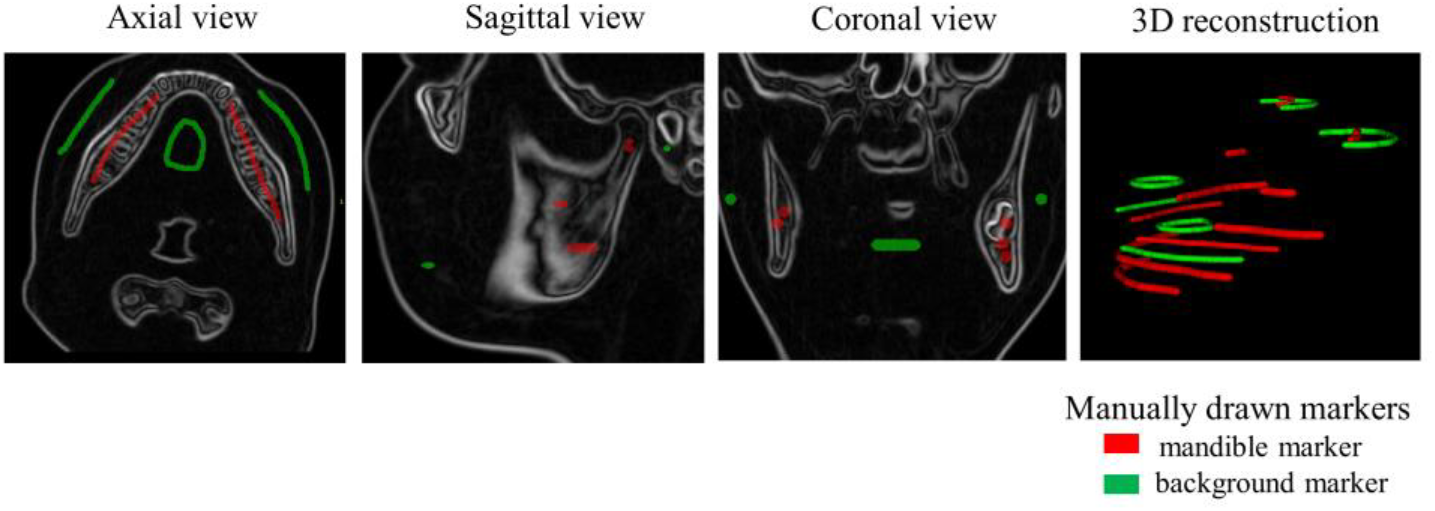
Manually drawn markers on the template image.

### Applying to a novel image

Markers were placed automatically inside and outside the mandible on a novel image from which the mandible was to be segmented. This was achieved by warping the template image onto each novel image using voxel-based image registration. Watershed markers, placed on the template, were warped along with this image, placing them into appropriate positions in each novel image.

In this study, the voxel-based image registration estimated a spatial transformation to be applied to each voxel of the novel image to corresponding voxels on the template image. The transformation involves both linear registration (translation and rotation) and non-linear registration (warp or stretch). The linear registration was used to coarsely align each test image to the template image using FLIRT registration in FSL open source tools (https://fsl.fmrib.ox.ac.uk/fsl/fslwiki/FSL). The non-linear registration then deformed the voxels on the test image more precisely into the template image using the advanced normalization tools, or ANTS (http://stnava.github.io/ANTs/). Further details on these methods are provided in the supplementary materials. Once the markers were automatically placed by the transformation, watershed segmentation proceeded as described above.

### Segmentation accuracy evaluation

Images of 20 adolescent subjects were used as test images in this study. The segmentation accuracy of the proposed method was assessed through the comparison to a well-accepted interactive segmentation method described in previous studies.^1,2^ This method was performed with open-source software ITK-SNAP (http://www.itksnap.org/pmwiki/pmwiki.php) by an experienced orthodontist (Hidden Content) and checked by a dentist (Hidden Content). Firstly, thresholding was used to grossly generate the main part of the mandible. The ‘region competition snake’ method was used to generate the condyles. Slice-by-slice editing in all three orthogonal views was required for further trimming the condyles and the lower teeth.

The segmented mandibles of these two approaches were compared by computing a Dice similarity coefficient for the overlapping voxels. This index ranged from 0 (no overlap) to 1 (complete overlap). The outer surfaces of the mandibles were generated with the marching cubes algorithm in MATLAB (https://au.mathworks.com/help/matlab/ref/isosurface.html). The boundary agreement between these two approaches were calculated as the surface distance between the two surfaces. This was quantified and visualized by a colormap.

## Results

### Timing

The interactive method typically required 30 to 40 minutes. Automatic segmentation of each mandible executed in 12-14 minutes on a Windows 7 PC running a Linux virtual machine.

### Accuracy

Mandibles segmented from the proposed automated method were compared against the interactive segmentation results. Dice similarity coefficients were 0.97 ± 0.01(mean ± SD), indicating almost complete overlap between the automatically segmented mandibles and the interactive segmented mandibles. Boundary deviations were predominantly under 1mm over most of the mandibular surfaces (Figure 3). The errors were mostly from the bones around partially erupted wisdom teeth, the condyles and the dental enamels, which had minimal impact on the overall morphology of the mandible (Figure 4).

**Figure 3.**
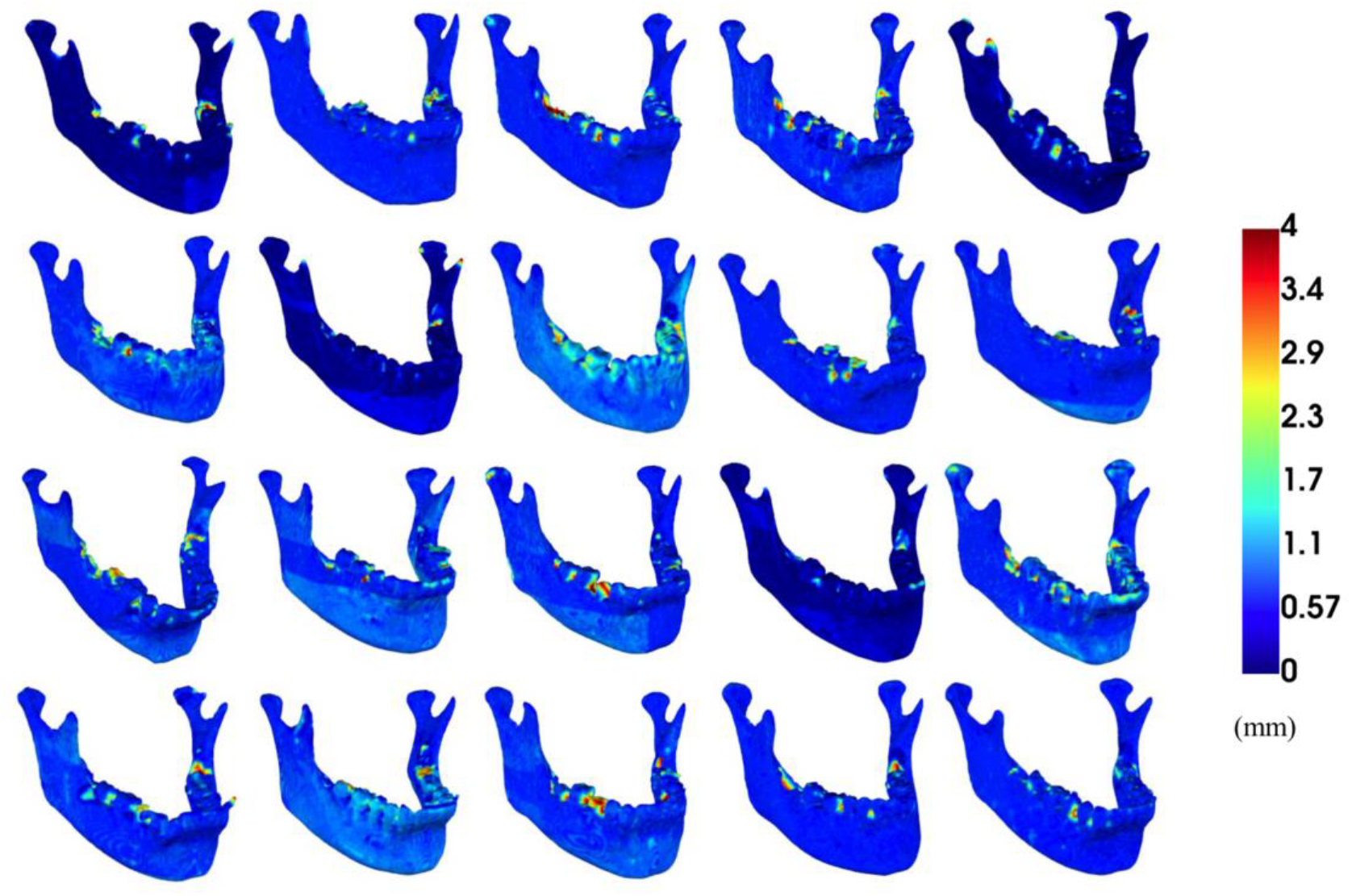
The discrepancy between the proposed automatic method and the interactive method in 20 test cases.

**Figure 4.**
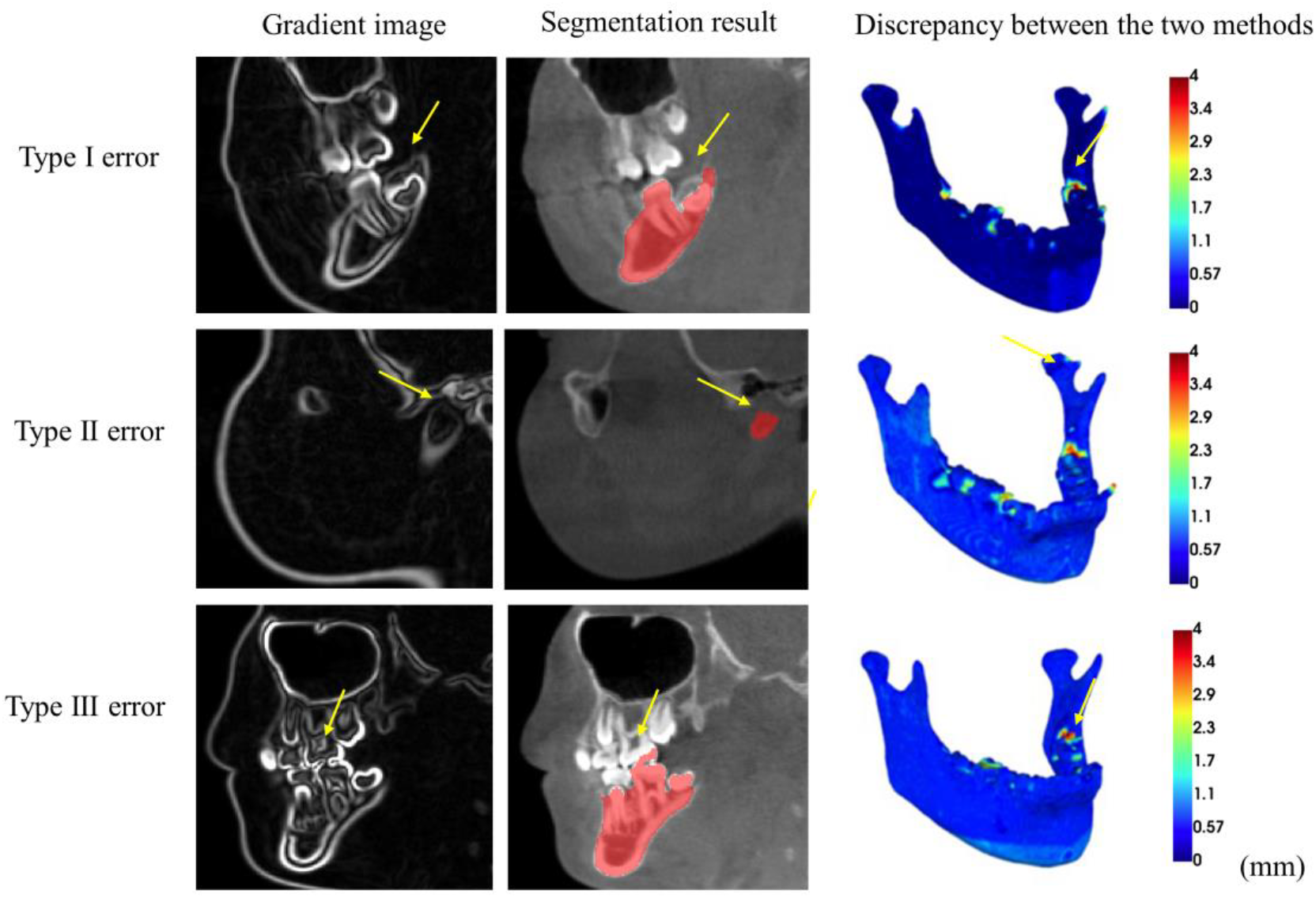
Segmentation errors for the proposed automatic approach. Type I error occurs at the partially erupted wisdom tooth. Type 2 error occurs at the ill-defined condyle. Type 3 error occurs at the dental enamel.

## Discussion

The quality of the mandibular segmentation determines the accuracy of subsequent applications, such as orthognathic treatment planning or orthodontic treatment evaluation. To date, most software-based mandibular segmentation involves continuous manual intervention, which is tedious and time-consuming, making it impractical for dealing with large numbers of subjects. In this study, we propose and evaluate an automated mandibular segmentation method using the marker-based watershed transform. This approach demonstrates time efficiency and comparable segmentation accuracy with a well-accepted interactive segmentation method.

In CBCT images, speckles or noise are more prominent than conventional CT images, which reduce the contrast resolution and make it difficult to differentiate low-density tissue in the image.^20^ The commonly provided algorithms in software show high sensitivity to image related artifacts which leads to reduced segmentation accuracy through inadequate structure capturing. For example, simple thresholding is effective in depicting the condyles from conventional CT images, where the image intensity histogram has a deep and sharp valley between two peaks representing the condyles and the soft tissue nearby. An adequate threshold can be chosen at the bottom of this valley to separate them apart. However, detecting the valley bottom precisely in CBCT images is difficult because the valley is flat and broad, imbued with noise. In this study, the Derivative of Gaussian filter is used to construct the height map, this not only enhances the intensity of the edges and dampen non-edges in the original image but also has noise suppression properties. Explicit placement of two markers, one inside and one outside the mandible ensures that the image is segmented into only two regions.

Another advantage of the proposed automated method is that it allows segmentation of the teeth. This is because the method is particularly useful for splitting touching objects, for example, it has been used to delineate touching cells or clustering nuclei from a microscopic image.^21^ In clinical situations, CBCT scans are often acquired with the upper and lower teeth touching, making them hard to separate using methods such as thresholding. The watershed approach is better able to separate touching teeth because the boundaries of the upper and lower teeth are accentuated in the gradient image, reflecting the sharp intensity changes between the enamels and air.

We have been able to fully automate the segmentation by automated watershed marker placement. This is achieved by aligning each test image to the template image using a voxel-based registration algorithm. Registration using a single template yielded good results for all the adolescent test cases in this study. However, human mandibles change markedly from infancy through childhood to adolescence, and from early adulthood to old age.^22^ Age-appropriate templates may be necessary for accurate image registration at different ages. This will ensure that the regions with high inter-age anatomical variability (such as the condyles and the coronoid processes) will be matched correctly and ensure the accuracy of the watershed marker placement.

Compared with the interactive segmentation, the automatic approach is time efficient and gives comparable accuracy. We demonstrate almost complete overlap between the automatically segmented mandibles and the interactively segmented mandibles in our test cases. There were, however, some errors at certain anatomical regions. First, the watershed flooding stops at the dental enamel of the partially erupted wisdom tooth before it reaches the cortical bone above as the intensity drop at the dental enamel is sharp. Second, the watershed lines are unpredictable at ill-defined condyles because of poor image quality for the cartilage in CBCT modality. Third, errors occasionally occur due to over-flooding to the enamels of the upper teeth. All these errors have minimal impact on the morphology of the mandible and can be easily fixed with a minimal amount of manual editing.

It should be noted that the interactive segmentation is less than a perfect gold standard. In this approach, a threshold was subjectively selected based on the intensity difference between the mandible and the rest of the structures. Differences in threshold selection due to blurring of the boundary or noise lead to slight changes in the final outline of the mandible. Slice-by-slice manual editing also results in jagged edges. The watershed method, on the other hand, implements a consistent definition of the boundary, corresponding to regions or rapid change in intensity. Therefore, some of the discrepancies between the interactively segmented mandible and the automated segmented mandible may be due to imperfections in the interactive segmentation.

## Conclusions

CBCT is increasingly used for diagnosis and treatment planning of the patients in implant dentistry, ENT, orthognathic surgery and interventional radiology. In this study, we propose and validate a practical and accurate marker-based watershed algorithm for automatically segmenting the mandible from low-dose CBCT images. Compared with user-depended interactive segmentation, our approach showed promising time efficiency and comparable accuracy. Further tests for images taken with different machine settings and from different age range patients are needed.

## Acknowledgements

The authors have no conflicts of interest to declare. We would like to thank (Hidden Content) for sharing his CBCT images; (Hidden Content) for providing the test cases and manually segmenting the mandibles. (Hidden Content) for the instruction in using ITK-SNAP. We also thank (Hidden Content) and (Hidden Content) for the valuable discussions.

